# Efficient Natural Plasmid Transformation of *Vibrio natriegens* Enables Zero-capital Molecular Biology

**DOI:** 10.1101/2023.08.11.553013

**Authors:** David A. Specht, Timothy J. Sheppard, Finn Kennedy, Sijin Li, Greeshma Gadikota, Buz Barstow

**Affiliations:** Cornell University, Biological and Environmental Engineering, Ithaca, NY 14853; Cornell University, Chemical and Biomolecular Engineering; Cornell University, Civil and Environmental Engineering

## Abstract

The fast-growing microbe *Vibrio natriegens* is capable of natural transformation where it draws DNA in from media via an active process under physiological conditions. Using an engineered strain with a genomic copy of the master competence regulator *tfoX* from *Vibrio cholera* in combination with a new minimal competence media (MCM) that uses acetate as an energy source, we demonstrate naturally competent cells which are created, transformed, and recovered entirely in the same media, without exchange or addition of new media. Cells are naturally competent to plasmids, recombination with linear DNA, and co-transformation of both to select for scarless and markerless genomic edits. The entire process is simple and inexpensive, requiring no capital equipment for an entirely room temperature process (Zero Capital protocol, 10^4^ cfu/*µ*g), or just an incubator (High Efficiency protocol, 10^5–6^ cfu/*µ*g). These cells retain their naturally competent state when frozen and are transformable immediately upon thawing like a typical chemical or electrochemical competent cell. Since the optimized transformation protocol requires only 50 minutes of hands-on time, and *V. natriegens* grows quickly even on plates, a transformation started at 9 AM yields abundant culturable single colonies by 5 PM. Further, because all stages of transformation occur in the same media, and the process can be arbitrarily scaled in volume, this natural competence strain and media could be ideal for automated directed evolution applications. As a result, naturally competent *V. natriegens* could compete with *E. coli* as an excellent chassis for low-cost and highly scalable synthetic biology.

## Introduction

Over the past decade, the fast-growing microbe *Vibrio natriegens* has attracted significant interest as the next generation replacement for *E. coli* as a host for synthetic biology and metabolic engineering.^1–5^ In addition to its extremely fast growth rate, with an optimal doubling time observed to be less than ten minutes in rich media, ^6^ *V. natriegens* has a number of advantages as a host for biotechnology, particularly for applications in metabolic engineering.^7^ Its fast growth rate is enabled by an extremely high rate of both rRNA production^8^ and substrate uptake, ^3^ even under anaerobic conditions. It is feedstock flexible, and in particular is capable of growth on low-energy substrates such as formate^9^ and acetate,^10^ and fixes nitrogen under anaerobic conditions.^11^ Like cloning strains of *E. coli*, it is BSL-1 and is genetically tractable, with a robust selection of promoters, terminators, and ribosomal binding sites,^12,13^ and has previously been engineered for production of numerous commercially-relevant compounds, including the bioplastic polyhydroxybutyrate (PHB),^14^ alanine,^3^ and 2,3-butanediol (2,3-BDO).^15,16^ Non-sterile seawater can be used as the source of water in growth media without substantial losses in yield, which has implications for sustainable bioproduction. ^16^ Additionally, *V. natriegens* appears to be capable of electron uptake from a cathode^17^ and is thus a potential chassis for electromicrobial production (EMP) in which electricity could be used directly to drive microbial metabolism and produce complex bioproducts from simple substrates (e.g. formate, acetate) or carbon fixed via engineered *in vivo* CO_2_ fixation pathways.^18,19^

While *V. natriegens* is a promising candidate for bioproduction, we contend that another major feature of biotechnological note is its extremely high potential for natural competence, in which cells can uptake intact extracellular DNA under specific environmental conditions. In many models of natural competence in gram negative microbes, for example in Mell et al.,^20^ natural transformation is often exclusively described as leading to linear ssDNA translocation across the inner membrane prior to homologous recombination. However, this does not appear to be the only possible fate of DNA taken up via natural competence, at least in *V. natriegens*. Natural transformation of intact plasmid DNA was first reported in a *Vibrio* in 1990,^21^ reported anecdotally in *V. natriegens* in Simpson et al.,^22^ and very recently contrasted with linear DNA uptake in a broad study of the genetic underpinnings of natural competency in *V. natriegens*.^23^

*Vibrionaceae* in general activate competence under starvation in the presence of chitin found in the shells of crustaceans.^24–26^ Artificial production of the master competence regulator *tfoX* allows competence to be induced in a more convenient manner, bypassing the need for chitin (but not starvation conditions). This is biotechnologically useful in part because it enables a simple means of conveying large, scarless genomic edits. In a technique called Multiplex Genome Editing by Natural Transformation (MuGENT), originally demonstrated in *Vibrio cholera*, high rates of cotransformation and subsequent homologous recombination of linear DNA enable the pairing of selectable and nonselectable genomic edits. ^27^ This was later extended to *V. natriegens* via expression of *tfoX* from *V. cholera*, where transformation rates were found to be as high as 1-10%.^14^ The method can be further enhanced by the deletion of genes which code for cytoplasmic nucleases which otherwise degrade transforming DNA. ^28^

While natural competence leading to homologous recombination has been a subject of attention of both biotechnology and fundamental biology, natural competence to dsDNA and plasmids generally, and in *V. natriegens* specifically, remains underexplored. This could be due in part to the existence of robust methods for creating chemically competent and electrocompetent cells in many microbes, including *V. natriegens*,^1^ which might obviate biotechnological interest in plasmid uptake via natural competence. In this article we make the case that natural plasmid transformation (NPT), which we treat distinctly from natural competence leading to homologous recombination, is biotechnologically valuable, especially for engineering at huge scales or in low-resource laboratory environments.

**Figure 1:**
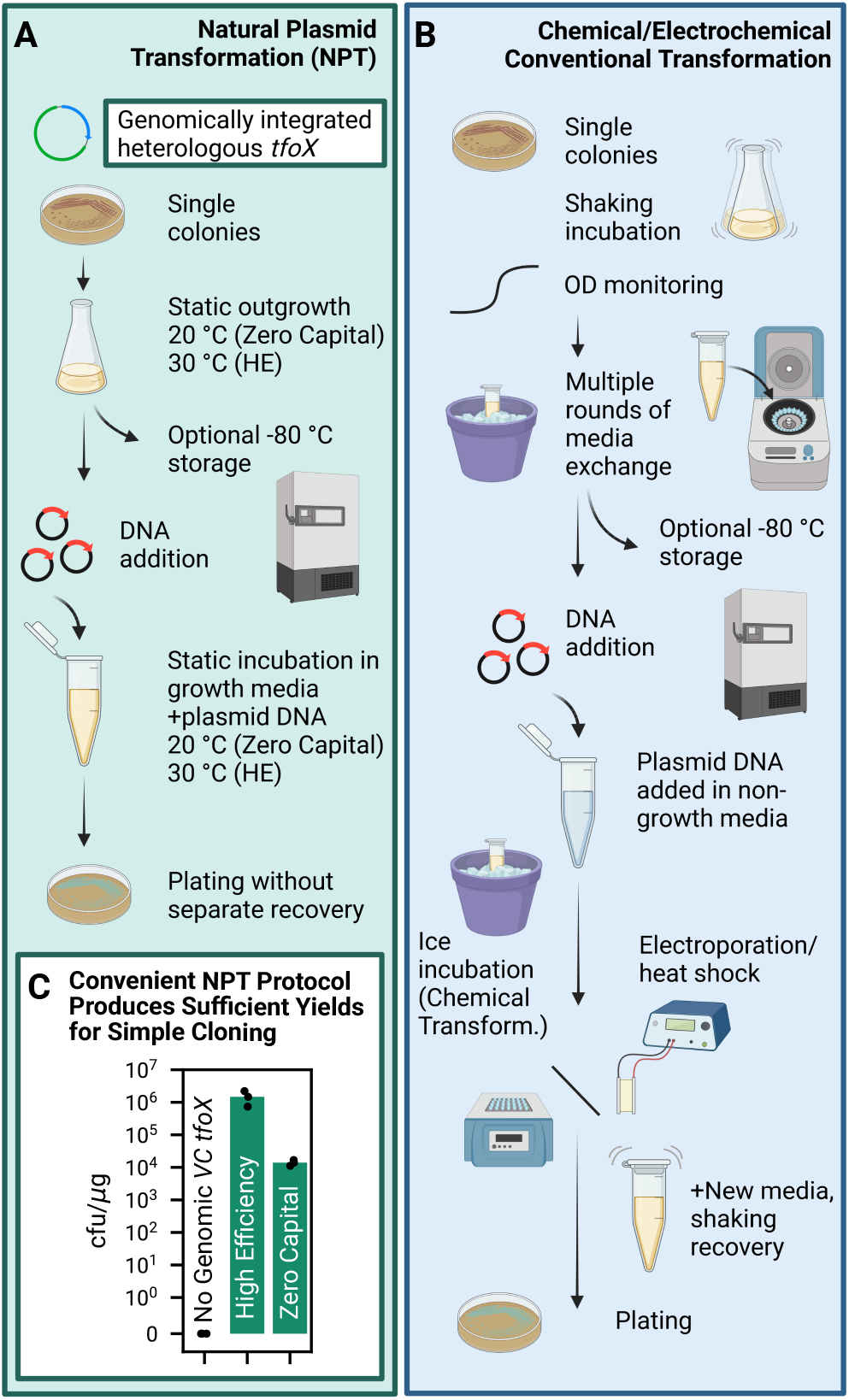
*Vibrio natriegens* genomically engineered for natural competence is transformable via direct addition of plasmid DNA to cells growing in a new minimal competence media (MCM). (**A**) Natural plasmid transformation (NPT) is enabled by genomic expression of *tfoX* from *V. cholera*, which allows for plasmid transformation without media exchange or addition, electroporation/heat shock, or a separate recovery step as is typical for conventional chemical or electrochemical competent cells (**B**). (**C**) We have developed two protocols, a High Efficiency protocol (HE, 10^6^ cfu/*µ*g), which requires only the use of an incubator and a deep freezer; and a Zero Capital protocol (10^4^ cfu/*µ*g), which requires no capital equipment at all and can be done entirely at room temperature. Because cells are transformed in their growth media, with no further concentration or media exchange, either protocol can be easily scaled, and the high efficiency transformation can be completed with as little as 50 minutes of hands-on time when started with frozen NPT competent cells.

In support of the biotechnological case for NPT, we produced a *V. natriegens* strain edited for natural competency, developed a pared-down media for growth, preservation, and transformation of cells, and originated an extremely simple protocol for creating and using these cells which optionally uses no capital equipment. Previous biotechnological usage of *V. cholera tfoX* expression in *V. natriegens* is predicated on use of a plasmid to express it.^14,29^ This presents a problem when using *V. natriegens* for plasmid cloning, and so we integrated *V. cholera tfoX* into the genome to obviate this issue. We invented a minimal competence media (MCM) which supports both growth and a state of natural competence which is maintained for 10s of hours. Cells are produced and transformed in this singular media, without any exchange, concentration, or separate media recovery, enabling the entire process to be able to be completed with no capital equipment at all, or enhanced with only the use of an incubator and deep freezer. Naturally-competent cells can be frozen and conveniently thawed for later use, to our knowledge for the first time by any researcher. We then show that NPT is practically useful, showcasing transformation of some typical cloning reactions, co-transformation of plasmid and linear PCR product for genomic editing, and then exhibit how the rapidity of both NPT and *V. natriegens* growth can be exploited to do transformation and then isolate single colonies by the end of a typical workday. We establish that natural competence is not just a biological curiosity or just a tool for genomic engineering, but allows for a useful third way of plasmid transformation, joining chemical and electrochemical competency.

Here, we report high efficiency plasmid uptake in an engineered *V. natriegens* which contains genomically-integrated expression of *tfoX* from *Vibrio cholera*,^14,24,29^ which we believe could be a useful chassis for molecular and synthetic biology research. Ultimately, we demonstrate that this strain can be used for efficient plasmid transformation, using a simplified shared media for competence expression, incubation, and recovery, enabling simple, scalable, low-capital plasmid engineering in *V. natriegens*.

## Results

### Genomic Integration of Heterologous *tfoX* Creates a *V. natriegens* Natural Competence Strain

Initial tests (not included) with *V. natriegens* containing either PMMB-tfoX (from Dalia et al.^14^) or pST 140 LVL2 cam (from Stukenberg et al.^29^) showed that ectopic expression of *V. cholera tfoX* via a plasmid can efficiently drive transformation of a second plasmid via natural transformation, as had been observed in Simpson et al.^22^ However, because these systems are predicated on plasmid expression of *V. cholera tfoX*, we sought to create a genomically-integrated version which could be used as a natural transformation-based host for molecular biology without interference from a second helper plasmid. Starting with the type strain (ATCC 14048), we used NT-CRISPR^29^ as described to perform a clean deletion of *dns*, which produces an extracellular nuclease which can reduce the efficacy of natural transformation^30^ and will likely degrade the quality of plasmid DNA. Subsequent editing was done in this Δ*dns* strain using a helper plasmid which expresses *tfoX* (pDS5.29) and the protocol for making genomic insertions via natural transformation as described in Dalia et al.^14^

Producing a strain with genomic expression of *tfoX* adequate to trigger measurable NPT proved to be unexpectedly difficult. Genomic insertion of two different versions of *tfoX* expression derived from pMMB-tfoX from Dalia et al.^14^ did not induce natural competence and two versions using the optimized genomic promoter P23 from Wu et al. ^13^ to drive *tfoX* were lethal. Ultimately, we observed one working version which came about spontaneously in a genomic insertion derived from pST 140 LVL2 cam^29^ which had been designed to contain *tfoX*, *camR*, and *lacI*. However, the working version which retained natural competence after curation of pDS5.29 contained a broken *lacI* sequence (IPTG inducibility of *tfoX* is lost) which we believe arose spontaneously during production of the tDNA template. Genomic sequences of both the region of interest and the entire engineered strain are included in the Supplementary Materials. Curiously, creation of a new strain with the same sequence but lacking *lacI* entirely did not exhibit natural competence, perhaps indicating a role of compositional context^31^ in the divergent promoter design used in Stukenberg et al.^29^ The working natural competence strain or its derivatives is used in all subsequent experiments.

### Development of Minimal Competence Media (MCM) Enables Preservation of Naturally Competent *V. natriegens* without Media Exchange

We next sought to reduce the protocol described in Dalia et al.^14^ such that both outgrowth and transformation could be accomplished in the same media, specifically optimizing for plasmid transformation. Briefly, in that protocol, cells are grown to saturation in a standard rich growth media (LBv2^1^). They are then diluted 1:100 in artificial seawater and incubated statically in the presence of tDNA for homologous recombination for 5 hours, prior to recovery and plating. As the original protocol is already quite simple, the original reasoning behind our work was to create a single media which could be used for many rounds of directed engineering without dilution in seawater. However, in the course of this research it was also discovered that in our minimal competence media (MCM), cells could be readily preserved in this state via flash-freezing and revived later for transformation. We sought to determine if these cells could be used as a drop-in replacement for traditional chemically or electrochemically competent cells, using an arbitrary plasmid with a pBR322 origin and GFP expression (pDS5.30).

MCM is comprised of a minimal set of essential ions and trace nutrients (Methods) with a low concentration of acetate being used as the carbon and energy source. During preliminary experiments, it was determined that the acetate concentration and pH were the predominant media determinants of the frequency at which NPT occurs. We did not find a substantial impact due to the concentration of any of the other media components, either in excess or in limiting conditions (including magnesium, nitrogen, phosphorus, and sulfur), nor was it beneficial to add any additional media components which are present at significant levels in natural seawater but inessential for growth (e.g. calcium). Transformation frequency is steady as a function of acetate concentration from 0.75 to 50 mM, but falls off at higher levels (Figure 2A). The total number of viable cfus are maximized at 3 mM acetate, and this is used for all subsequent experiments. Similarly, a sodium concentration of 350 mM is used in all subsequent experiments as this maximizes the total number of viable cells (Figure 2B). The natural transformation is highly sensitive to pH, however, as transformation becomes undetectable below pH 6.5 (Figure 2C), despite the fact that there are a sufficient number of untransformed cells to detect a transformation frequency as low as 10^-6^ (Figure S1). pH is adjusted to 7.4 and buffered using HEPES to center it in the optimal range for all subsequent experiments.

**Figure 2:**
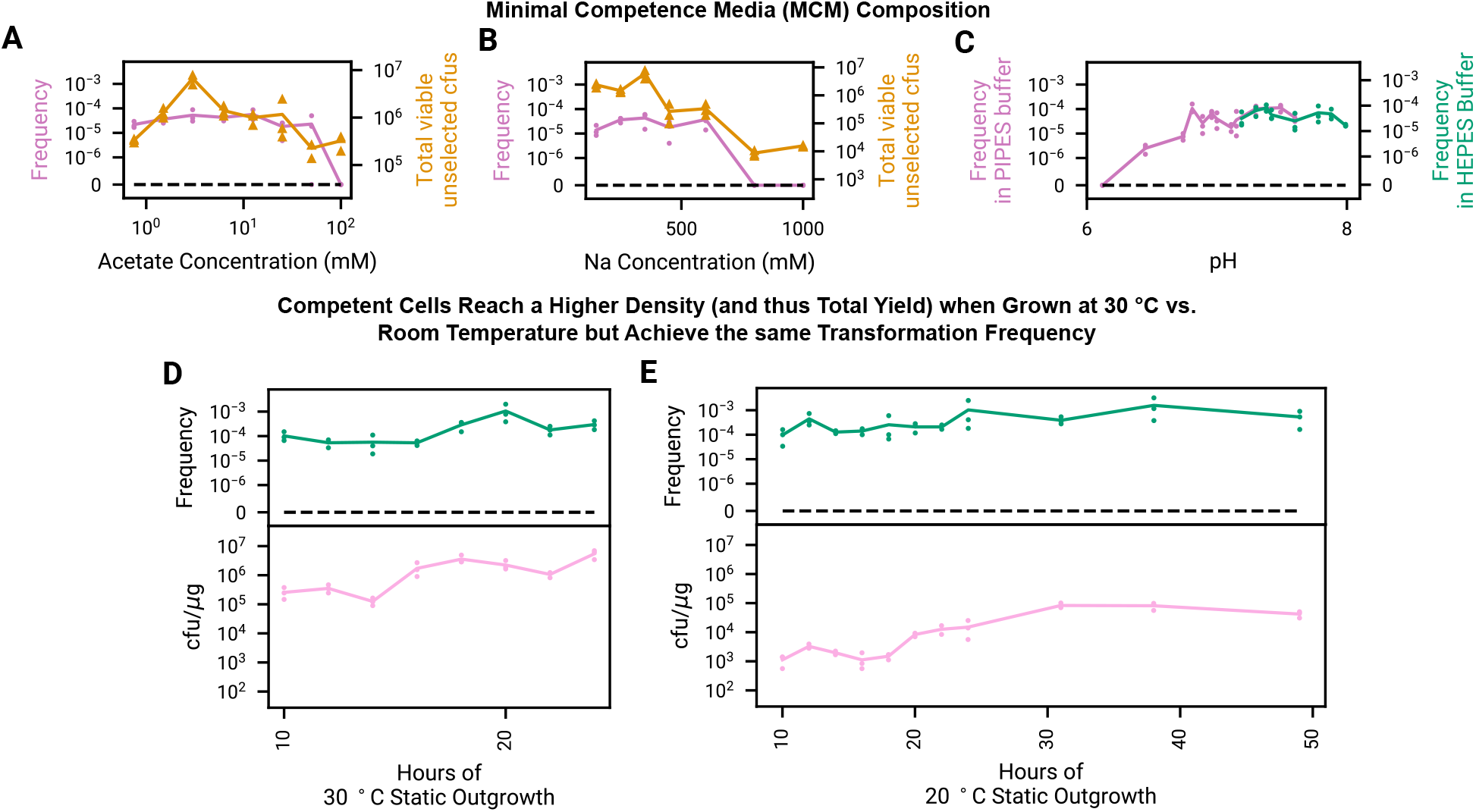
Minimal competence media (MCM) derived from essential seawater components plus acetate is used to both grow out and freeze *V. natriegens* in a state of natural competence. (**A**,**B**) Transformation frequency (transformed cfus/untransformed viable cfus) is insensitive to acetate and sodium concentration over a wide range of concentrations, but overall growth (number of viable cfus in 50 *µ*L) is maximized close to 3 mM acetate and 350 mM sodium. (**C**) Transformation is pH dependent and robust in pH ranging from 7 to 8, but falls to zero at lower pH despite growth at pH as low as 6 sufficient to detect natural transformation (Figure S1). In order to center pH in an area with robust transformation and cfu yield, MCM is used at a final pH of 7.4, buffered by HEPES, for all subsequent experiments. (**D**,**E**) Transformation frequency is relatively insensitive to the period of time that cells are grown out statically prior to transformation, but the overall number of transformed cfus increases as the cells grow in MCM, and cells achieve a much higher maximal density under 30 *^◦^*C growth. In all of the above figures, cells are flash frozen in glycerol after outgrowth, thawed, incubated statically with 25 ng of DNA for 30 minutes at 30 *^◦^*C for transformation, and recovered in LBv2 with shaking at 37 *^◦^*C for 1 hour.

### Natural Plasmid Transformation (NPT) is Streamlined to Create an Extremely Simple Transformation Protocol with Minimal Hands-on and Total Runtime

Briefly, single colonies are used to inoculate 20 mL of MCM, which is grown statically overnight for a period of time at either 30 or 20 *^◦^*C. After this outgrowth period, 350 *µ*L aliquots are taken, mixed with glycerol, and then flash frozen prior to storage and then subsequent transformation. To transform, cells are thawed, plasmid DNA is added, and cells are incubated statically with the plasmid DNA. In earlier experiments (Figure 2), cells are recovered at 37 *^◦^*C in a shaking incubator in rich media for 1 hour and then plated.

Transformation is sensitive to the length and temperature of the outgrowth time, with frequency and cfu/*µ*g yield maximized around 18-20 hours for outgrowth at 30 *^◦^*C (Figure 2D) and around 24-40 hours at 20 *^◦^*C (Figure 2E). For the case of room temperature outgrowth, the state of natural competence is maintained for at least over a day after the maximal cell density is reached, without substantial loss in transformation frequency or yield after 50 hours of outgrowth. In subsequent experiments, cells are collected at 18 hours when grown at 30 *^◦^*C, and at 24 hours when grown at 20 *^◦^*C. Shaking cells during this initial outgrowth stage destroys subsequent competence.

We next sought to optimize the protocol for transforming the cells (Figure 3), finding that because cells remain metabolically active during NPT many aspects of traditional transformation protocols can be omitted. In order to make a transformation protocol which is competitive with chemical transformation in terms of hands-on time, we sought to limit the transformation time to approximately 1.5 hours (which would be typical for a chemical transformation with a 30 minute incubation and 1 hour of recovery, neglecting the other steps). Given a time budget of 1.5 hours, we were surprised to find that a 30 minute static incubation followed by addition of a recovery media (LBv2, Recovery Media,^1^ or MCM) under typical recovery conditions (shaking at 37 *^◦^*C) performs more poorly than simply incubating cells statically for 1.5 hours with no recovery step at all and plating cells immediately after incubation (Figure 3A). Cells are insensitive to being melted at room temperature (Figure 3B) and are robust and can be vortexed for up to 60 seconds at maximum speed prior to the addition of plasmid DNA (Figure 3C). Shaking during incubation reduces transformation frequency by over an order of magnitude but does not destroy it (Figure 3D).

**Figure 3:**
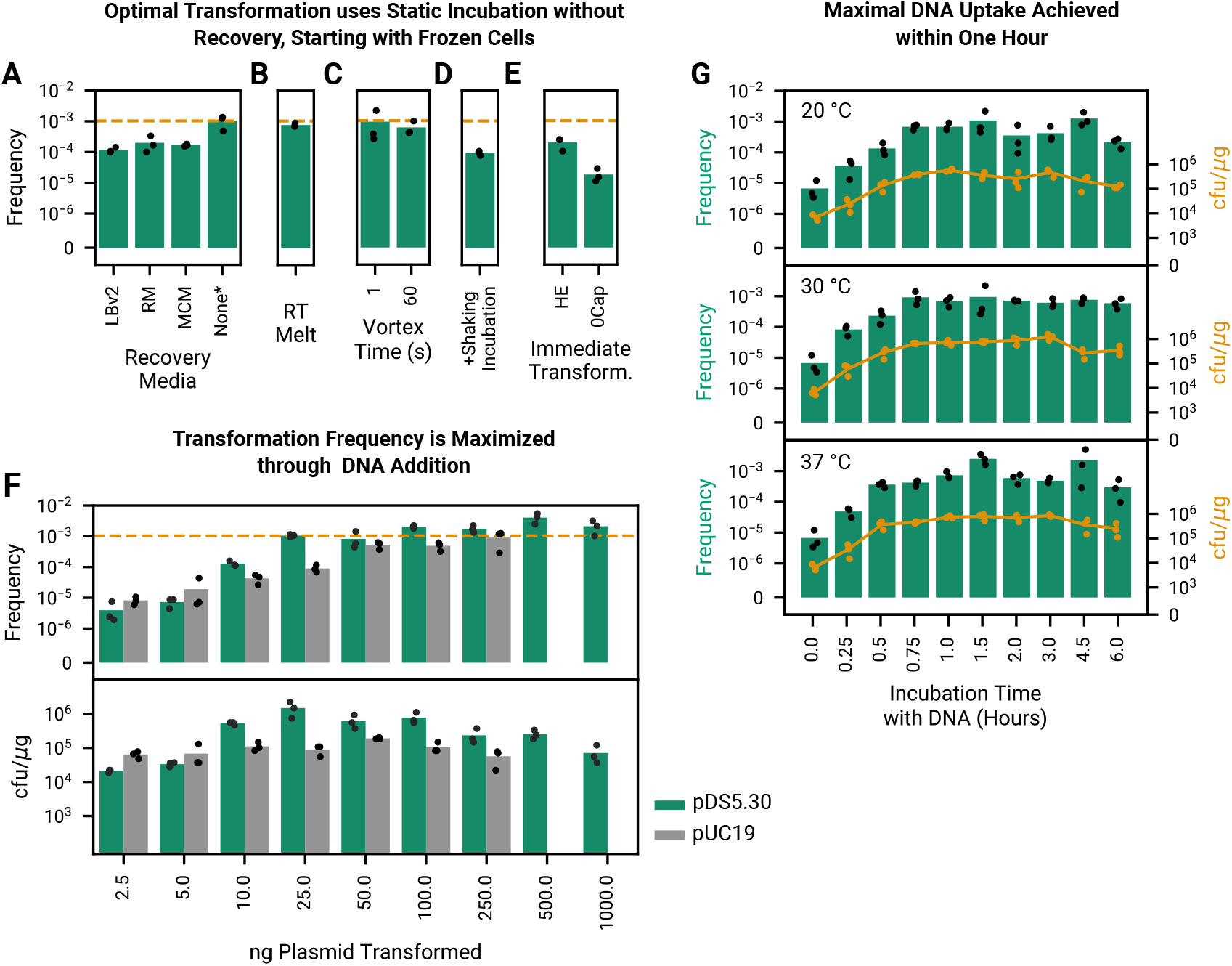
Optimization of transformation of naturally competent cells shows that natural plasmid transformation (NPT) is extremely fast and remains robust even if many details of working with traditional competent cells are neglected. (**A**) Given a limit of 90 minutes of hands-on time, allowing cells to incubate for 90 minutes in MCM yields a higher transformation frequency than a 30 minute incubation followed by addition of fresh media (LBv2, recovery media (RM),^1^ or MCM) and 60 minutes of shaking incubation. Because cells are metabolically functional during the active transformation process, they immediately begin to express antibiotic resistance genes, eliminating the value of a separate recovery step. The performance of subsequent experiments are compared to the no recovery condition (dashed orange line). (**B,C**) Cells can be thawed at room temperature (RT Melt) with no loss in efficiency, and prior to DNA addition can even be vortexed at full speed for up to a minute with minimal losses. (**D**) Cells lose an order of magnitude in transformation efficiency if shaken during incubation rather than being left to take up and express DNA statically. **E**) We consider full immediate transformation without flash freezing at 30 and 20 *^◦^*C (starting with cells grown out in 30 or 20 *^◦^*C, respectively, and proceeding directly after the prior outgrowth step to transformation), corresponding to our High Efficiency (HE) and Zero Capital (0Cap) protocols. Interestingly, cells consistently lose approximately an order of magnitude in transformation frequency if they are not flash frozen after they are created. (**F**) In 350 *µ*L of unconcentrated cells in media, yield (transformed cfu per *µ*g of added plasmid DNA) is optimized around 25 ng of added pDS5.30 DNA. Transformation yield with pUC19 is lower. (**G**) In all temperature conditions, optimal transformation efficiency and yield is reached within 45 minutes of incubation. While eventual transformation yield is sensitive to initial growth temperatures (Figure 2D,E), incubation with plasmid DNA is accomplished efficiently and comparably from 20-37 *^◦^*C in a window up to 3 hours, with overall yield falling off slightly after that period. 25 ng of plasmid DNA is used for experiments in all subplots except F.

We next tested the impact of using cells immediately after overnight outgrowth, rather than after -80 *^◦^*C storage, and found that freezing cells enhances transformation efficiency by almost an order of magnitude (Figure 3E). We grew out, incubated, and plated each experiment entirely at either 30 or 20 *^◦^*C. The 20 *^◦^*C protocol requires no capital equipment at all (no OD meter, centrifuge, incubator, shaker, freezer, ice machine, heat bath/electroporator, or deep freezer is required), and is the basis of our Zero Capital Protocol (Note S3), while the 30 *^◦^*C protocol only requires an incubator. The addition of glycerol prior to flash-freezing is not the driver of increased transformation frequency, as the addition of glycerol has no impact when cells are immediately transformed after outgrowth (Figure S2).

We optimized the transformation frequency and yield (cfu per *µ*g of added transforming DNA) as a function of the amount of added DNA (Figure 3F) using both pDS5.30 and an arbitrary plasmid commonly used to report transformation efficiency commercially (pUC19). Transformation frequency increases rapidly as up to 25 nanograms of DNA are added to a standard 460 *µ*L aliquot of cells, with marginal increases in total frequency as up to several hundred nanograms are added, with cfu/*µ*g yield maximized around 50-250 nanograms. pDS5.30 consistently provides a higher yield at these concentrations.

Transformation frequency and yield is insensitive to incubation temperatures ranging from 20 to 37 *^◦^*C, increasing with incubation time until plateauing around 45 minutes (Figure 3G). Transformation in which cells are plated immediately after DNA addition with no incubation at all results in a surprisingly high transformation frequency (*≈* 10^-5^), perhaps indicating that some of this activity may be occurring on the plate as dilutions dry. Incubation on ice results in no transformants (not shown). This result, in combination with the prior result finding that transformation efficiency is not improved by the addition of recovery media, indicates that cells begin expressing antibiotic resistance genes from the plasmid immediately upon uptake during static incubation in MCM, unlike with traditional chemically competent cells which are inactive when incubated on ice.

High energy food sources like sugars inhibit NPT in a singular media where there is no subsequent dilution or media exchange step. Using the minimal MCM components, and a diverse array of potential metabolites in lieu of acetate at the same 3 mM concentration (glucose, sucrose, gluconate, formate, pyruvate, sorbitol, and tryptone), we sought to see if there might be options other than acetate as a carbon/energy source in MCM. We found that pyruvate is the next-best inducer of natural transformation (Figure S3), which is competent at a frequency on the order of 10^-4^. A mixture of 10% LBv2 and 90% artificial seawater also exhibited transformation, although at a low frequency on the order of 10^-6^. Replication of the creation of competent cells using pyruvate MCM and 10% LBv2 with flash freezing and -80 *^◦^*C storage shows improved transformation frequency (Figure S4), as was the case for acetate MCM.

We tested the use of our finalized protocol optimized for our genomically-integrated expression of *tfoX* on the Δ*dns* strain containing the plasmid pMMB-tfoX^14^ in which we did some of our initial observations of NPT. IPTG induction of *tfoX* from was lethal to cells grown in MCM, while cells without the addition of IPTG showed normal rates of survival, potentially indicating some toxic effect resulting from excessive *tfoX* overexpression in combination with the MCM environmental condition.

### Naturally Competent *V. natriegens* is an Effective Tool for Plasmid Cloning

Transformation of frozen competent cells through isolation of single colonies is possible within a standard workday due to the combination of rapid *V. natriegens* growth and the fact that a 45 minute incubation is sufficient to maximize NPT transformation frequency and yield (Figure 4G). As a simple demonstration of this capability, we began a transformation at 9 AM (Figure 4A). NPT competent cells were removed from the deep freezer and then thawed on the benchtop for *≈* 5 minutes. Transforming plasmid DNA was then added and cells were incubated for 45 minutes at 30 *^◦^*C, and then immediately plated (37 *^◦^*C) on pre-warmed LBv2 agar plates using plating beads at around 10 AM. By 3:30 PM, extremely small single colonies were visible, which we then imaged at 4:30 PM and used a single one to inoculated a culture tube of LBv2. By 9 AM on the following day, that overnight culture yielded an OD of 9.2 in 3 mL of media.

**Figure 4:**
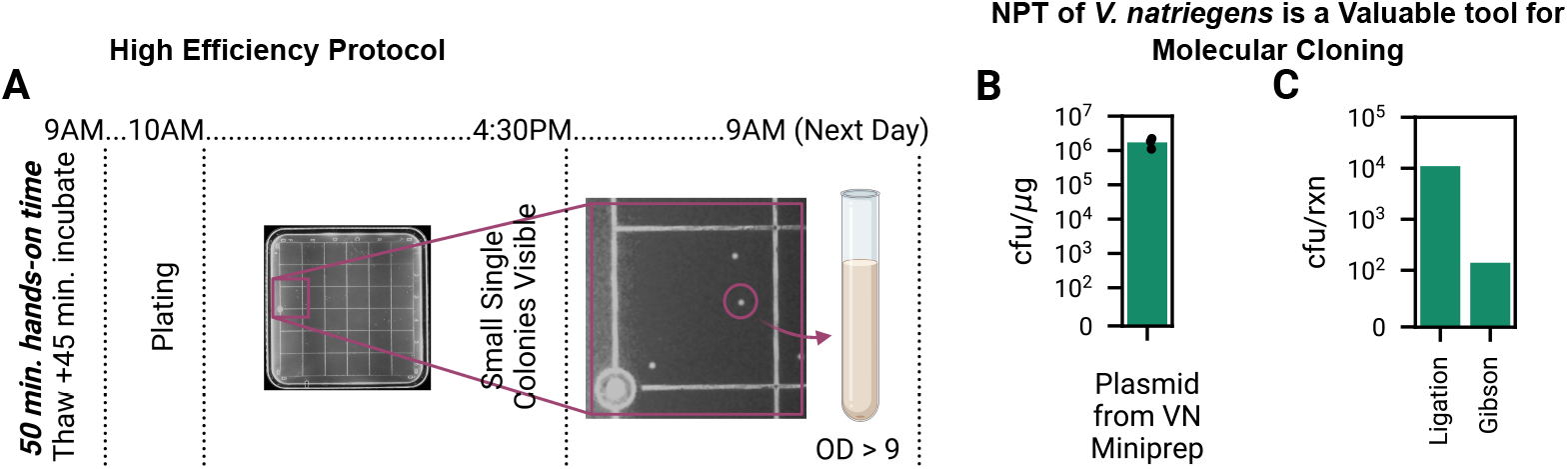
NPT (Natural Plasmid Transformation) of *V. natriegens* produces culturable single colonies within a standard workday, empowering rapid cloning. (**A**) In our rapid, high efficiency protocol (requiring only an incubator and deep freezer), *V. natriegens* transformation and growth is so fast that single colonies are visible (*≈* 0.7 mm) and can be picked within a standard workday when transformation is done first thing in the morning. (**B**) Plasmids miniprepped from *V. natriegens* are transformable back into *V. natriegens* with the same yield as those produced from *E. coli* DH5*α*. (**C**) Arbitrary, common molecular biology reactions (a ligation via Kinase, Ligase, and Dpn1 (KLD), deleting GFP expression from pDS5.30; and a Gibson assembly, inserting GFP expression into a pUC19 backbone) can be transformed directly into the natural competence strain of *V. natriegens* without any additional cleaning steps, yielding 11,000 and 140 transformants, respectively, from 2 *µ*L of reaction product.

The ability to get to the single colony stage within a standard workday is significant and gives a putative cloning strain of *V. natriegens* an edge in utility over *E. coli*. Upon plating, transformants of *E. coli* cloning strains will not be visible to the naked eye after 6 hours of growth. As a result, outgrowth of colonies is typically carried out overnight. This means that, depending on timing, many protocols which typically take two days to complete can be done in one if started in the morning.

Plasmids can be readily extracted from *V. natriegens* using standard miniprep kits developed for use with *E. coli*. These plasmids can then be used to transform *V. natriegens* with the same yield as plasmids derived from *E. coli* DH5*α* (Figure 4B).

We next demonstrated that two arbitrary molecular biology reactions can be carried out in *V. natriegens* NPT competent cells (Figure 4C). We used KLD, an enzyme mixture of kinase, ligase, and dpn1 optimized to ligate PCR product and remove confounding template plasmid DNA in one reaction, to delete GFP expression from pDS5.30, with a yield of 11,000 transformed cfus from a 2 *µ*L reaction (the standard volume recommended by NEB for a transformation with *E. coli*). We also used Gibson assembly to add GFP expression to pUC19, with a yield of 140 transformed cfus from a 2 *µ*L reaction.

### Co-transformation of Plasmid and Linear DNA Enables Scarless Genomic Editing

Because natural transformation is a highly non-Poisson process such that an uptake event is strongly correlated with additional DNA uptake,^32^ it should be possible to co-transform both selectable plasmid DNA and arbitrary linear DNA at a high rate. The objective of this is to pair unselected genomic edits with an arbitrary plasmid transformation (Figure 5A), similarly to how MuGENT simultaneously pairs selectable and nonselectable genomic edits.^14,27^ In all previous experiments, the natural competence strain of *V. natriegens* still contained the chloramphenicol resistance gene *camR*. This is unnecessary and undesirable for subsequent cloning which might utilize chloramphenicol, and thus we sought to delete it using the plasmid/linear DNA co-transformation. Using 400 ng of transforming DNA with 3 kB homology arms, paired with 25 ng of pDS5.30, we co-transformed these and determined that 22% of cells received the desired edit (8 out of 36 colonies tested for loss of chloramphenicol resistance). Cells are then easily cured of pDS5.30 and we verified that the resulting strain had the expected sequence and remained naturally competent, although with reduced transformation frequency (Figure 5B). The reduction in transformation frequency may relate to the fine tuning of *tfoX* expression and sensitivity to compositional context,^31^ as was the case in sensitivity to proximal LacI expression discussed previously.

**Figure 5:**
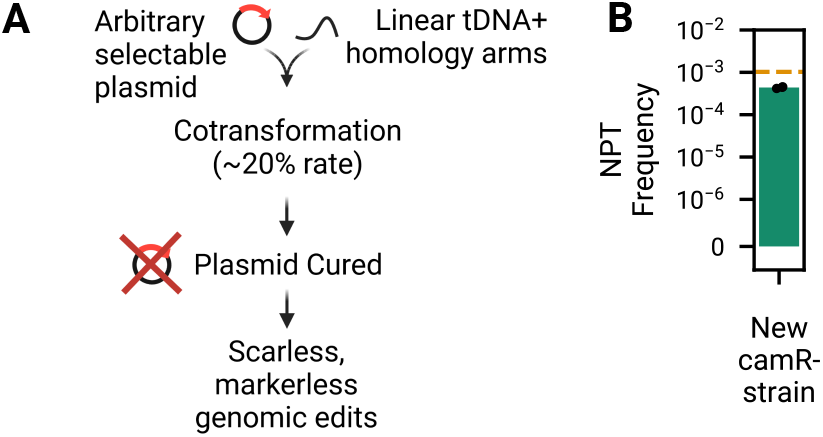
Cotransformation of plasmids and linear DNA enables rapid scarless and markerless genomic edits of *V. natriegens*. (**A**) As in MuGENT,^28^ unselected genomic edits can be paired with selectable markers due to a high rate of natural competence co-transformation. By using a plasmid to convey selection, the resulting strain can be easily cured of the plasmid without leaving a selectable marker behind in the genome. (**B**) We use this co-transformation to remove *camR* from the strain used throughout this study and the resulting strain retains natural competence.

## Discussion

In this work, we have shown that a genomically engineered strain of *V. natriegens* created specifically for enhanced natural competence can be used as an effective chassis for low cost and low capital plasmid engineering. Natural plasmid transformation (NPT), using our strain and specific transformation media, allows for alternate growth and high-efficiency transformation without exchange of media. Further, cells can be flash frozen and stored in this naturally competent state. Plasmids can be co-transformed with linear DNA for genomic editing, enabling diverse scarless genomic edits without requiring additional genomic insertions ^28^ or additional plasmid engineering for CRISPR counter-selection.^29^ Due to the low resource intensity and yet high efficiency of the transformation protocol, this work supports the idea that *V. natriegens* is a strong candidate next-generation molecular biology workhorse, especially for low-resource laboratory environments. In particular, we believe that a chassis which readily grows and can be edited entirely at room temperature, using no capital equipment, could be a radical new resource for the democratization of synthetic biology, especially in education.

NPT is a curious third alternative to chemical or electrochemical transformation of plasmids in *V. natriegens*. Many microbes are capable of a state of natural competence under the right conditions,^33^ and many more are likely to be as specific environmental triggers are discovered. Even *E. coli*, the current molecular biology workhorse, is capable of natural transformation of plasmids while on solid media^34,35^ independent of the type IV pilus,^36^ although the efficiency and protocol make it impractical for routine cloning. In general, natural competence in bacteria is controlled by an eclectic set of diverse environmental and physiological conditions. While natural transformation is used extensively as a tool to modify diverse microbial genomes, few microbes have previously been engineered specifically for enhanced natural competence, most notably *B. subtilis* ^37,38^ and *V. cholera*.^27^ *B. subtilis* is a poor substitute for *E. coli* due to its evolutionary distance, preference for multimeric plasmids, and poor diversity of plasmids mutually compatible with *E. coli*.^39^ To our knowledge, there are no demonstrations of cells in a naturally competent state with comparable general utility to traditional frozen *E. coli* -based chemical competent cells.

The uptake of intact circular, double-stranded DNA runs counter to models of natural transformation in gram negative microbes which are predicated on solely ssDNA translocation across the inner membrane prior to homologous recombination, e.g. as described in Mell et al.^20^ Retraction of type IV competence pili which bind dsDNA draws it to the cell surface and mediates DNA internalization, driving natural competence.^40^ dsDNA could then be taken up by putative membrane elements which pass dsDNA through the inner membranes, as appears to be the case for limited natural transformation of *E. coli* on solid media.^41^ Alternatively, it is also possible that two ssDNA plasmid monomers are taken up individually and reconstituted inside the cell, as is the case in *S. pneumoniae*.^42^ Transformation of plasmid monomers could be used to differentiate between such single- and double-hit kinetics, as is done in Sun et al.^43^ The fact that our system depends on *tfoX* expression (Figure 1C), which is an essential trigger for type IV pilus-based DNA uptake and homologous recombination,^14,27^ indicates an effective comingling of systems for both double and single-stranded DNA uptake. Very recent work^23^ identifies a new membrane protein which appears to be essential for genomic linear tDNA integration but not plasmid DNA uptake. A complete knockout collection would be useful to understand the underlying molecular machinery for dsDNA translocation across the inner membrane as well as enhancing NPT for biotechnological use, for example by enhancing NPT in rich media or during rapid growth. Our observation that the frequency of transformation is significantly enhanced by flash freezing also indicates that there could be some role of physical trauma to the cell membrane which either promotes competence gene expression or directly facilitates DNA uptake.

Expression of *V. cholera tfoX* may need to cross some threshold in order to trigger natural competence. However, excess expression of *tfoX* under competence inducing environmental conditions can be lethal, as we observed in our initial efforts to engineer genomic overexpression. This reflects a need to appropriately balance the genomic trigger for natural competence with environmental conditions. The fact that deletion of the entire faulty *lacI* construct also destroys natural competence, as discussed in the Results, demonstrates a sensitivity to the nearby genomic context. We suspect that the tightly packed divergent^31^ PLacI and Ptac promoters in the original design from Stukenberg et al.^29^ may play a role in upregulating *tfoX* in such a way that it promotes natural competence even when genomically integrated, although this has not been explored further.

NPT can be efficiently accomplished with 50 minutes of hands-on time, less than half of the time required for chemically competent cells (Figure 4A), and requires neither a heat shock nor electroporation. Additionally, NPT competent cell preparation is extremely simple, requiring no wash steps and functioning within a wide time window which does not require monitoring of OD (Figure 2D). It is a dynamic process by which cells actively take up DNA in a physiologically-controlled process at temperatures relevant for growth, as opposed to chemical transformation which occurs via diffusion while cells are maintained on ice. The fact that cells remain metabolically active under these conditions likely explains why a separate recovery step is not required in order to express antibiotic resistance genes (Figure 3A). In total, this simplicity is automation friendly and enables some unique applications. For example, it would be possible to independently transform collections of many clonally-isolated individuals, at room temperature and with no separate chilling or shaking steps, in 96-well plates in MCM.

The extremely high rate of natural transformation in *V. natriegens* with *tfoX* expression makes NPT practically relevant in routine cloning applications (Figure 4B,C). However, as a matter of yield in transformed cfu per added microgram of DNA, it is not yet competitive with commercial strains of cloning *E. coli*. Yields are lower than practically similar one-step chemical transformation methods in *E. coli*, which also do not require media exchange or heat shock and achieve a transformation efficiency of *≈* 10^7^ - 10^8^ cfu/*µ*g.^44^ However, because natural transformation is an active process by which cells take up DNA under metabolically functional conditions, it is likely that the rate of natural transformation can be increased through engineering as has been done previously in *V. cholera*,^27^ either by increasing the frequency of competence pili production or retraction, or by making the process insensitive to growth in rich media. Due to their fragility, traditional competent cells are shipped on dry ice and are stored in -80 *^◦^*C deep freezers. Researchers have attempted to create commercial versions of dehydrated^45^ and freeze-dried^46^ chemically competent cells, but to our knowledge no version has ever gone on the market. We speculate that naturally competent cells with a genomically enhanced capability for transformation could be used to create commercially-relevant fridge or shelf-stable competent cells via dehydration or freeze drying.

Aside from general cloning usage, the fact that cells can be grown, transformed, and subsequently recovered in the same media all at the same temperature without media exchange could have powerful advantages in continuous evolution. For example, in MAGE (Multiplexed Automated Genome Engineering),^47^ DNA for recombineering is delivered via electroporation and the process is thus bottlenecked by transformation efficiency.^48^ Especially given the feedstock flexibility and extremely high growth rate of *V. natriegens*, NPT of nonreplicative plasmids and directed evolution could represent a potent combination. While *E. coli* can also be grown and transformed in a shared media,^44^ cells must be chilled during transformation and recovered under growth conditions prior to antibiotic selection.

Other novel applications are possible. Recently published work^49^ demonstrates DNA detection via natural competence in *B. subtilis*, which is an excellent chassis due both to its high degree of natural competence as well as the possibility of using spores in an eventual application. However, a freeze-dried *V. natriegens* could work just as well and simultaneously benefit from rapid growth for speedier sequence detection.

In its simplest manifestation (Figure 1A), our protocol demonstrates that NPT of *V. natriegens* can be accomplished with minimal capital resources. Since *V. natriegens* can be dried and shipped on filter paper, a putative end user could revive the cells, make them competent via our protocol, and transform them using no capital equipment at all, save for what is required to make sterile media. Either way, no fridge, freezer (-80 *^◦^*C or conventional), OD meter, centrifuge, heat block, incubator, electroporator, or shaker is required as part of the transformation process, conducted entirely at room temperature. Such a transformation could be a useful tool outside of traditional biology lab environments, useful for diverse new users including high school educators, iGEM teams, or physics or chemistry laboratories seeking to do some limited synthetic biology.

Ultimately, one must ask the question of whether researchers would actually switch to a replacement for *E. coli* for molecular cloning. While *V. natriegens* has a multitude of advantages and could serve as a drop-in substitute in many applications, for example in protein production^50^ and simple DNA assembly (Figure 4C), dominance of classic *E. coli* strains like DH5*α* is due to their proven reliability over decades. However, the reliance on the hardware of a textbook biology lab limits the reach of synthetic biology. NPT via *V. natriegens* which enables low-resource end users to use the tools of molecular biology will push the field forward and further the democratization of synthetic biology.

## Materials & Methods

### Working with *V. natriegens*

In general, growth of *V. natriegens* was done in LBv2 liquid media or LBv2 agar plates.^1^ When necessary, antibiotics were used in both solid and liquid culture at a final concentration of 200 *µ*g/mL (kanamycin), 2 *µ*g/mL (chloramphenicol), and 10 *µ*g/mL (carbenicillin, see Note S1). In instances where it was necessary to transform *V. natriegens* via conventional means, cells were made electrocompetent using the protocol as described in Weinstock et al.^1^ Glycerol stocks were created by mixing cells in late exponential growth (*≈*OD 1) at a ratio of 3:1 with 60% glycerol prior to storage in a -80 *^◦^*C freezer. In instances where artificial seawater media was used, we filter sterilized 28 g/L of Instant Ocean Sea Salt in deionized water.

### Genomic Editing of *V. natriegens*

A *dns* knockout of the *V. natriegens* wild type strain (ATCC 14048) was created using NT-CRISPR, using the protocol as described in Stukenberg et al.^29^ to do the insertion as described in Dalia et al.,^14^ followed by the CRISPR-based counterselection. The requisite 3kB homology arms were assembled in pUC19 via Gibson assembly (NEBuilder HiFi DNA Assembly), creating pDS5.13, from which tDNA was amplified. Counterselection was accomplished using the spG Cas9 NT-CRISPR plasmid (pST 140 LVL2 cam, Addgene 179334) with spacer sequence tgcactatccagtgccgccg (pDS5.17).

We then inserted the *tfoX/lacI/camR* construct from Stukenberg et al.^29^ into the Δ*dns V. natriegens*. In order to accomplish this, we created a second helper plasmid (pDS5.29) containing *tfoX* and GFP to aid in eventual curation. Using this helper plasmid and tDNA derived from pDS5.27 (containing the insertion and homology arms), we inserted the construct into the genome following the protocol for natural transformation from Dalia et al.^14^ Cells were plated for chloramphenicol resistance, and verification of the insertion at the expected location was confirmed by colony PCR. As in NT-CRISPR, cells were then cured of the helper plasmid via 37 *^◦^*C antibiotic-free growth in LBv2 for 6 hours, plating a 10^-7^ dilution on LBv2 agar plates, and then striking colonies onto kanamycin plates in order to verify helper plasmid loss.

### NPT Protocol Development

In all iterations, cells of the natural competence strain or the negative control were first struck out from a glycerol stock onto LBv2 plates for single colonies. On the subsequent day, a single colony was used to inoculate 20 mL of media in 100 mL flasks to create cells with a state of natural competence. In instances where cells are preserved in the -80 *^◦^*C freezer, 350 *µ*L of overnight culture is added to 110 *µ*L of 60% glycerol, mixed by pipetting up and down, placed on ice for several minutes, and then flash frozen in liquid nitrogen. In early iterations, cells from the freezer are thawed on ice (Figures 2, 3A), while in subsequent experiments cells are thawed on the benchtop. In instances where cells are used for immediate transformation without freezing, DNA is directly added to 350 *µ*L of overnight culture, except in Figure S2 where 110 *µ*L of 60% glycerol is also added. Except in cases where the amount of DNA is explicitly changed (Figure 3F), 25 ng of plasmid DNA is used for all NPT transformations.

Cells are then incubated in the presence of the tDNA. This is done statically, with the exception of Figure 3D, for a period of time ranging from 0 to 6 hours, at either 30 °C or room temperature. In early iterations, (Figure 2), 1 mL of LBv2 is added and cells are placed in a shaker at 37 *^◦^*C for recovery. After it became clear that there was limited benefit from the addition of recovery media (Figure 3A, discussed in the main text), cells are plated immediately after incubation, diluting as appropriate in order to calculate transformation efficiencies. Plated cells are grown out at 37 *^◦^*C, except for in Figure 3E 20 *^◦^*C which is a demonstration of the fully room temperature protocol.

In its final manifestation, MCM consists of: 9 mM HEPES, 3 mM sodium acetate, 1.9 mM ammonium chloride, 1.6 mM potassium phosphate, 7 mM potassium chloride, 1 mM magnesium sulfate, 31 mM magnesium chloride, and 350 mM sodium chloride. In order to prevent precipitation, 1 mL of undilute hydrochloric acid is used to lower the pH of 900 mL of deionized water prior to adding the media components and water to a total volume of 1 L. The final mixture is then adjusted upwards to pH 7.4 using 1 M sodium hydroxide and sterile filtered. The media will precipitate if autoclaved. This recipe is used for all experiments in included figures except Figures 2A-C, indicated in pink, where 10 mM PIPES is used in lieu of HEPES and the pH is adjusted to 7, and in Figures S3 and S4, where various carbon/energy compounds are used in lieu of acetate, as indicated.

We have written instructions for NPT in our natural competence strain, describing both the high speed/efficiency and low capital transformations, as protocols in Note S2 and Note S3, respectively.

With the exception of Figure 3F where indicated in gray, all plasmid transformations were done using pDS5.30, a plasmid with a pBR322 origin which expresses kanamycin resistance and GFP.

### Development of NPT as a Tool for Cloning

In order to test the utility of NPT and *V. natriegens* as a host for cloning, we designed two arbitrary Gibson assembly and KLD reactions, creating final plasmids pDS5.43 and pDS5.44, respectively. The requisite PCRs were completed using NEB Hot Start Q5. PCR product for Gibson assembly was cleaned using a Zymo Clean & Concentrator kit and then used in a NEBuilder HiFi DNA Assembly reaction as described by the manufacturer. In the KLD reaction (NEB KLD Enzyme Mix), the PCR product is used directly in the KLD reaction without further cleaning. In both, 2 *µ*L of reaction product is added directly to the competent cells in media, just as with plasmid DNA in the previously described NPT protocol.

Miniprep extraction of plasmid DNA from *V. natriegens* was accomplished using the E.Z.N.A Plasmid DNA Mini Kit I produced by Omega Bio-Tek, following the manufacturer’s instructions.

For the demonstration of producing single colonies within a standard workday, colonies are imaged using an Azure Biosystems Gel Imaging System. The image contrast is altered in order to highlight the presence of colonies and facilitate measurement of their size.

### Co-transformation of Plasmid DNA with Linear tDNA for Genomic Editing

tDNA with 3 kB homology arms (as described in Dalia et al.^14^), designed for the deletion of *camR* which was previously inserted, was created by assembly of plasmid pDS5.45 via Gibson assembly, from which tDNA was amplified with PCR. 25 ng of pDS5.30 was co-transformed with 400 ng of tDNA. Co-transformation of plasmid DNA and tDNA for genomic editing was done using the described protocol for plasmid transformation in our natural competence strain. Once plated, we arbitrarily struck 36 colonies onto chloramphenicol plates in order to estimate the frequency of genomic editing. Deletion of *camR* was additionally verified by colony PCR. The remaining inserted sequence was then Sanger sequenced.

## Acknowledgements

This work was supported by a Cornell Energy Systems Institute Postdoctoral Fellowship to D.A.S., ARPA-E Grant DE-AR0001608 (Advancing a Low Carbon Built Environment with Inherent Utilization of Waste Concrete and CO2 via Integrated Electrochemical, Chemical, and Biological Routes, ADVENT), a Cornell 2030 Project Fast Grant, and a generous gift from Mary Fernando Conrad and Tony Conrad to B.B. Sequencing was done with resources provided by the BRC Genomics Facility (RRID:SCR 021727) at the Cornell Institute of Biotechnology. All plasmid sequences used in this work are available and hosted by Benchling. We would like to thank Ankur Dalia for his gift of plasmid PMMB-tfoX and Anke Becker for her gift of pST 140 LVL2 cam (Addgene plasmid #179334).

## Attribution

Conceptualization, D.A.S and B.B.; Investigation, D.A.S, T.J.S., and F.K.; Writing - Original Draft, D.A.S; Writing - Review & Editing, D.A.S., T.J.S, and B.B.; Funding Acquisition, S.L., G.G., and B.B.; Resources, B.B.; Supervision, B.B., G.G.

