## Supplementary Materials for "Efficient Natural Plasmid Transformation of *Vibrio natriegens* Enables Zero-capital Molecular Biology"

David A. Specht,† Timothy J. Sheppard,† Finn Kennedy,† Sijin Li,‡ Greeshma Gadikota,¶ and Buz Barstow*,†

†*Cornell University, Biological and Environmental Engineering, Ithaca, NY 14853*

‡*Cornell University, Chemical and Biomolecular Engineering, Ithaca, NY 14853*

¶*Cornell University, Civil and Environmental Engineering, Ithaca, NY 14853*

#### Note S1 Ampicillin vs. carbenicillin sensitivity in *V. natriegens*

While ampicillin and carbenicillin are treated interchangeably when used for *E. coli* selection, we have observed that in *V. natriegens* carbenicillin exhibits dramatically stronger selection on solid media than ampicillin does. In our tests, concentrations as low as 2 $\mu$g/mL of carbenicillin can be sufficient for counterselection, while there are colonies which escape ampicillin selection at concentrations as high as 50 $\mu$g/mL in freshly made plates. Ultimately, we use carbenicillin at a concentration of 10 $\mu$g/mL for selection of pUC19 recipients (Methods). A prior review of methods in *V. natriegens* antibiotic concentrations[5] shows that there may not have been contrasting usage of these two antibiotics in prior studies.

### Note S2 NPT protocol optimized for high efficiency

1. Prepare: LBv2 plates[1] with and without the appropriate antibiotic; 60% sterile glycerol; liquid nitrogen (if flash freezing);

2. Prepare 1× minimal competence media (MCM): 9 mM HEPES, 3 mM sodium acetate, 1.9 mM ammonium chloride, 1.6mM potassium phosphate, 7 mM potassium chloride, 1 mM magnesium sulfate, 31 mM magnesium chloride, and 350 mM sodium chloride. In order to prevent precipitation, 1 mL of undilute hydrochloric acid is used to lower the pH of 900 mL of deionized water prior to adding the media components and water to a total volume of 1 L. The final mixture is then gradually adjusted upwards to pH 7.4 using 1 M sodium hydroxide and sterile filtered. The media will precipitate if autoclaved.

3. Strike cells from a glycerol stock onto LBv2 plates for single colonies. Incubate overnight at 37 °C.

4. From a single colony, inoculate 20 mL of MCM in a sterile flask. Incubate for 18 hours, statically, at 30 °C.

5. Briefly resuspend cells by swishing the flask, and take as many 350 $\mu$L aliquots as needed. These can either be added to 110 $\mu$L 60% glycerol, flash frozen, and stored at -80 °C, or used immediately in subsequent steps.

6. If using frozen cells, thaw them at room temperature for $\approx$ 5 minutes.

7. Add $\geq$ 25 ng of plasmid DNA to the cells. Invert or vortex briefly to mix.

8. Allow cells to incubate statically for at least 45 minutes at 37 °C. Per Figure 3G, cells can be incubated for up to 3 hours with minimal additional gains to transformation frequency or yield at temperatures ranging from 20 to 37 °C.

9. Dilute in MCM as necessary to get single colonies (for a typical transformation, 1-2 orders of magnitude is sufficient), spread onto a prewarmed LBv2 plate, and grow at 37 °C for single colonies. Small colonies are visible 6-7 hours after plating. Efficiency is reduced if cells are diluted in a medium other than MCM.

### Note S3 Room temperature NPT protocol optimized for no capital equipment

1. Prepare: LBv2 plates[1] with and without the appropriate antibiotic; 60% sterile glycerol.

2. Prepare 1× minimal competence media (MCM): 9 mM HEPES, 3 mM sodium acetate, 1.9 mM ammonium chloride, 1.6 mM potassium phosphate, 7 mM potassium chloride, 1 mM magnesium sulfate, 31 mM magnesium chloride, and 350 mM sodium chloride. In order to prevent precipitation, 1 mL of undilute hydrochloric acid is used to lower the pH of 900 mL of deionized water prior to adding the media components and water to a total volume of 1 L. The final mixture is then gradually adjusted upwards to pH 7.4 using 1 M sodium hydroxide and sterile filtered. The media will precipitate if autoclaved.

3. Strike cells from a glycerol stock onto LBv2 plates for single colonies. From this point on, no capital equipment is necessary. When grown at room temperature, single colonies will become visible 24 hours after being struck out.

4. From a single colony, inoculate 20 mL of MCM in a sterile flask. Incubate statically at room temperature for 24 hours (although competency is maintained for at least up to 50, Main Figure 2E).

5. Briefly resuspend cells by swishing the flask, and take as many 350 $\mu$L aliquots as needed.

6. Add $\geq$ 25 ng of plasmid DNA to the cells. Invert to mix.

7. Allow cells to incubate statically for at least 45 minutes at room temperature. Spread on a room temperature LBv2 plate. Small colonies are visible 24 hours after plating at room temperature.

### Supplementary Figures

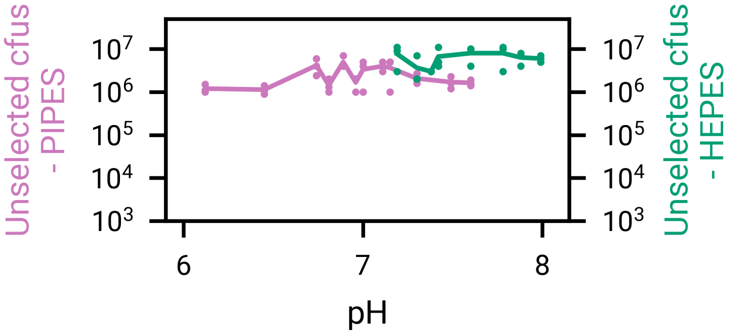


Supplementary Figure S1: **Total survivorship during transformation is slightly increased under higher pH in HEPES buffer.** Total number of unselected colony forming units in 50 $\mu$L of MCM after transformation, as a function of pH, buffered with either PIPES or HEPES, as indicated. *V. natriegens* readily grows in MCM from at least pH 6.12 to 7.99. Despite this, NPT is pH-dependent (Figure 2C), occurring at a reduced frequency for lower pH, and is undetectable at pH 6.12.

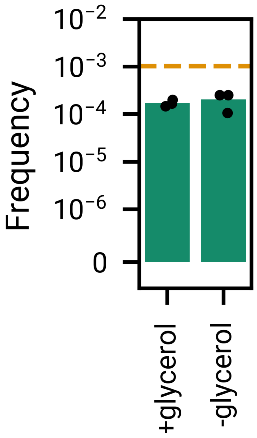


Supplementary Figure S2: **The addition of glycerol in immediate transformation of cells after 30 °C outgrowth does not restore the transformation frequency observed in the case of flash freezing.** This indicates that the addition of glycerol is not the driver of increased transformation frequency.

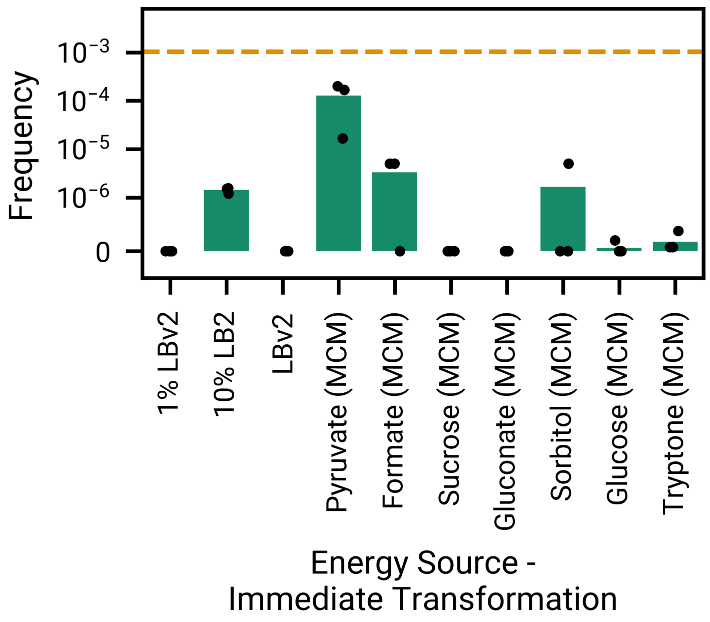


Supplementary Figure S3: **Pyruvate is an effective alternative carbon/energy source in lieu of acetate.** Under immediate transformation after outgrowth (with no flash freezing and -80 °C storage), pyruvate is the next-best carbon/energy source in lieu of acetate for driving natural transformation using MCM. All carbon/energy sources in MCM are at 3 mM. Additionally, a mixture of 10% LBv2 and 90% Instant Ocean Media produces measurable transformation.

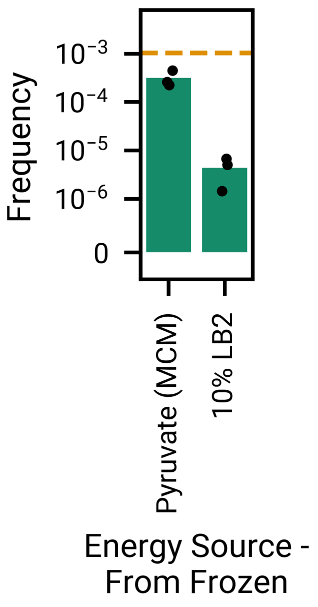


Supplementary Figure S4: **Pyruvate MCM and 10% LBv2 can be frozen and preserved in a natural competence state.** As in the primary experiments with acetate-based MCM, freezing and -80 °C storage prior to transformation enhances transformation in pyruvate MCM and 10% LBv2 (compare with Figure S3).
